## Supplemental Materials for "Dysregulated *H19*/*Igf2* expression disrupts cardiac-placental axis during development of Silver Russell Syndrome-like mouse models"

**Supplemental Table 1. Primers and PCR conditions utilized in this study.**

| Primer | Forward | Reverse | Reference |
|---|---|---|---|
| **Mouse qRT-PCR primers** | | | |
| *Arbp* | TCCCACTTACTGAAAAGGTCAAG | TCCGACTCTTCCTTTGCTTC | (Hur et al., 2016) |
| *Nono* | GCTCGTGAGAAGCTGGAGAT | TTCTTGACGTCTCATCAAATCC | (Plasschaert & Bartolomei, 2014) |
| *Rpl13a* | ATCCCTCCACCCTATGACAA | GCCCCAGGTAAGCAAACTT | |
| *H19* | GTCTCGAAGAGCTCGGACTG | ACTGGCAGGCACATCCAC | (Hur et al., 2016) |
| *Igf2* | CGCTTCAGTTTGTCTGTTCG | GCAGCACTCTTCCACGATG | (Weaver et al., 2009) |
| **Mouse allele-specific expression primers** | | | |
| *Igf2* | ATCTGTGACCTCTTGAGCAGG | GGGTTGTTTAGAGCCAATCAA | (de Waal et al., 2015) |
| PCR condition: 95°C 2min, (95°C 15s, 58°C 10s, 72°C 20s)x26-30 cycles, 72°C 5min | | | |
| **Mouse genotyping primers** | | | |
| *hIC1* | CCTTCACGGCTTTGACACTC | GTCAACCGGAGGCACAGTAT | (Hur et al., 2016) |
| *⊿H19* | TTGTGGTGAGGCTGTCTTTG | CCTATTCCCCATTCCATCCT | This study |
| *⊿3.8* | CCAACTGAGAGGGCCATAGTGTGAG | CCACAGAGTCAGCATCCAC | (Thorvaldsen et al., 2002) |
| PCR condition: 95°C 2min, (95°C 15s, 58°C 10s, 72°C 20s)x35 cycles, 72°C 2min | | | |
| **gRNA pairs for generating *⊿H19* allele** (PAM sequences are underlined) | | | |
| Pair A | CTTCAATATAATGCGACTCATGG | AACGTGCGCTGGAACGATACAGG | |
| Pair B | CAATATAATGCGACTCATGGGGG | ATCAGTACATGGCCCCGCCGGGG | |
| Primers for px335 plasmid amplification | TTAATACGACTCACTATAGGNNNNNNNNNNNNNNNNNNNNNGTTTTAGAGCTAGAAATAGC (The underlined nucleotides were substituted with each gRNA sequence excluding the PAM sequence) | AGCACCGACTCGGTGCCACT | |

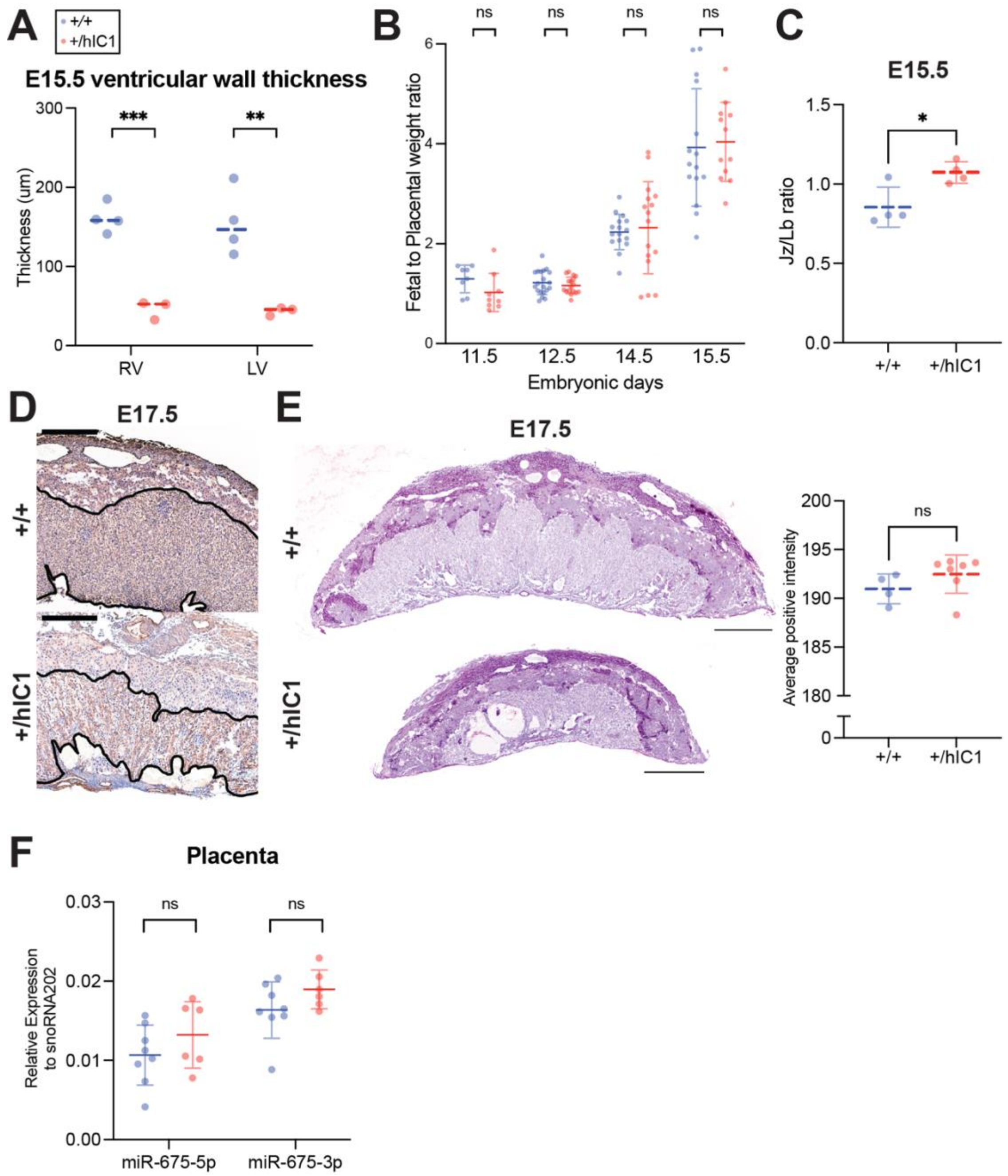


**Supplemental Figure 1. Supplementary data for anomalies observed in +/*hIC1* placentas.**

(A) Quantification of ventricular wall thickness (µm), measured from E15.5 wild-type and *+/hIC1* hearts (mean ± SD). 4 wild-type and 3 *+/hIC1* embryos from two different litters were examined. (B) Fetal to placental weight ratios of the wild-type (blue) and *+/hIC1* (red) samples at E11.5, E12.5, E14.5 and E15.5. Each data point represents an average ratio of each genotype from one litter. (C) Junctional zone (Jz) to labyrinth (Lb) ratio in E17.5 wild-type and *+/hIC1* placentas. (D) Example of labyrinth zones that were used to quantify the microvessel density in CD34 immunostained E17.5 wild-type and *+/hIC1* placental sections from Figure 3G. Thrombi in *+/hIC1* placentas were excluded. (E) (Left) Representative images of

PAS staining counterstained with hematoxylin on E17.5 wild-type and *+/hIC1* male placental sections. (Right) Quantification of PAS stained images. (F) Relative total expression of miR-675-3p and miR-675-5p in E15.5 wild-type and *+/hIC1* placentas (mean ± SD). (A, C, E, F) Each data point represents an individual conceptus from different litters. Scale bars = 1mm. Statistics used are (A, C, E, F) multiple unpaired t-tests, (B) multiple paired t-test. *$P < 0.05$, **$P < 0.01$, ***$P < 0.001$, ****$P < 0.0001$, ns = not significant.

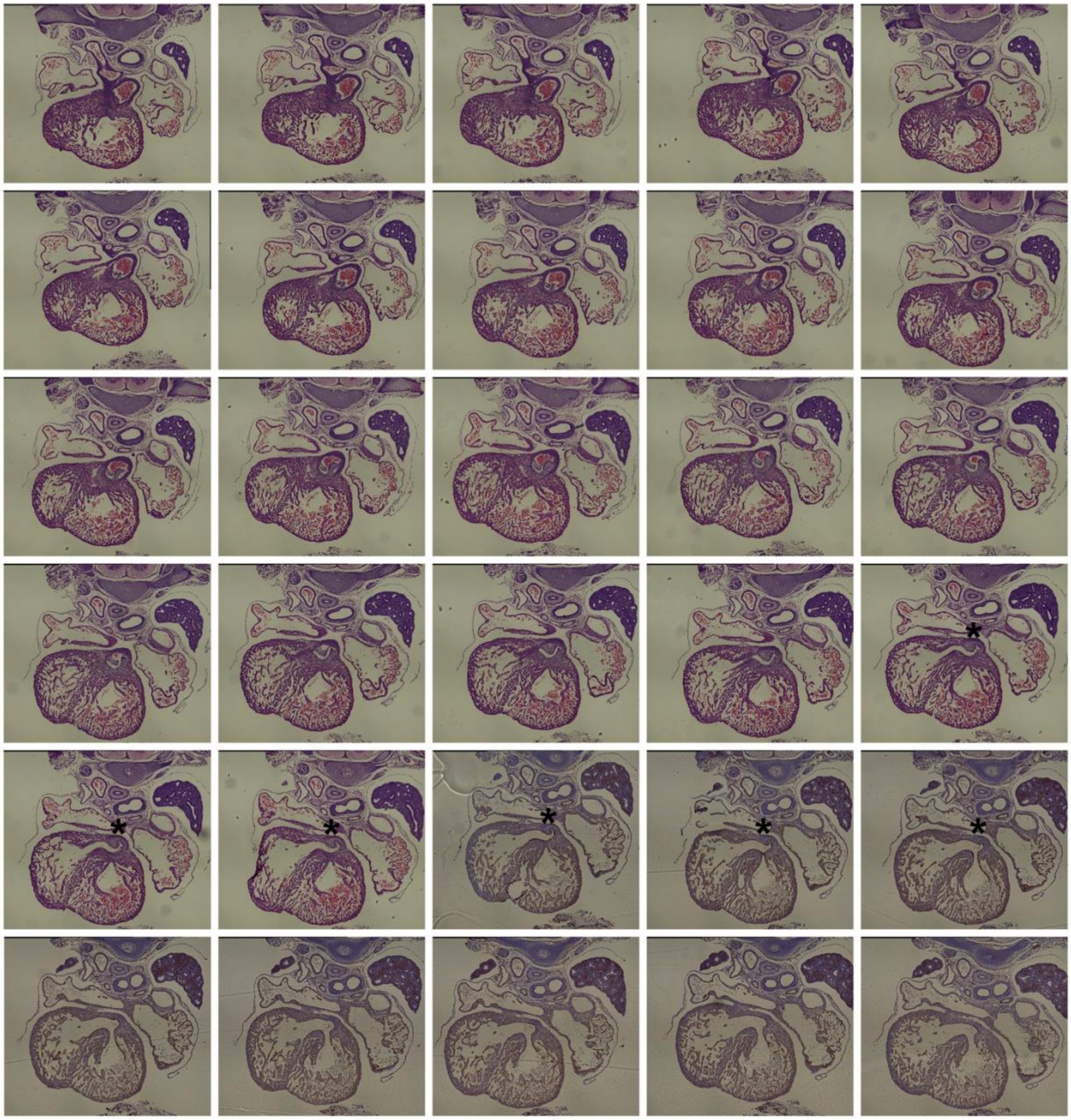

**Supplemental Figure 2. Serial cross-sections of E15.5 *+/hIC1* embryonic heart with BPV.**

Sequential histological sections demonstrating the bicuspid pulmonary valve (BPV) phenotype. Rudimentary anterior cusp can be seen fused with the right pulmonary cusp (raphe) in sections denoted by * above the pulmonary valve.

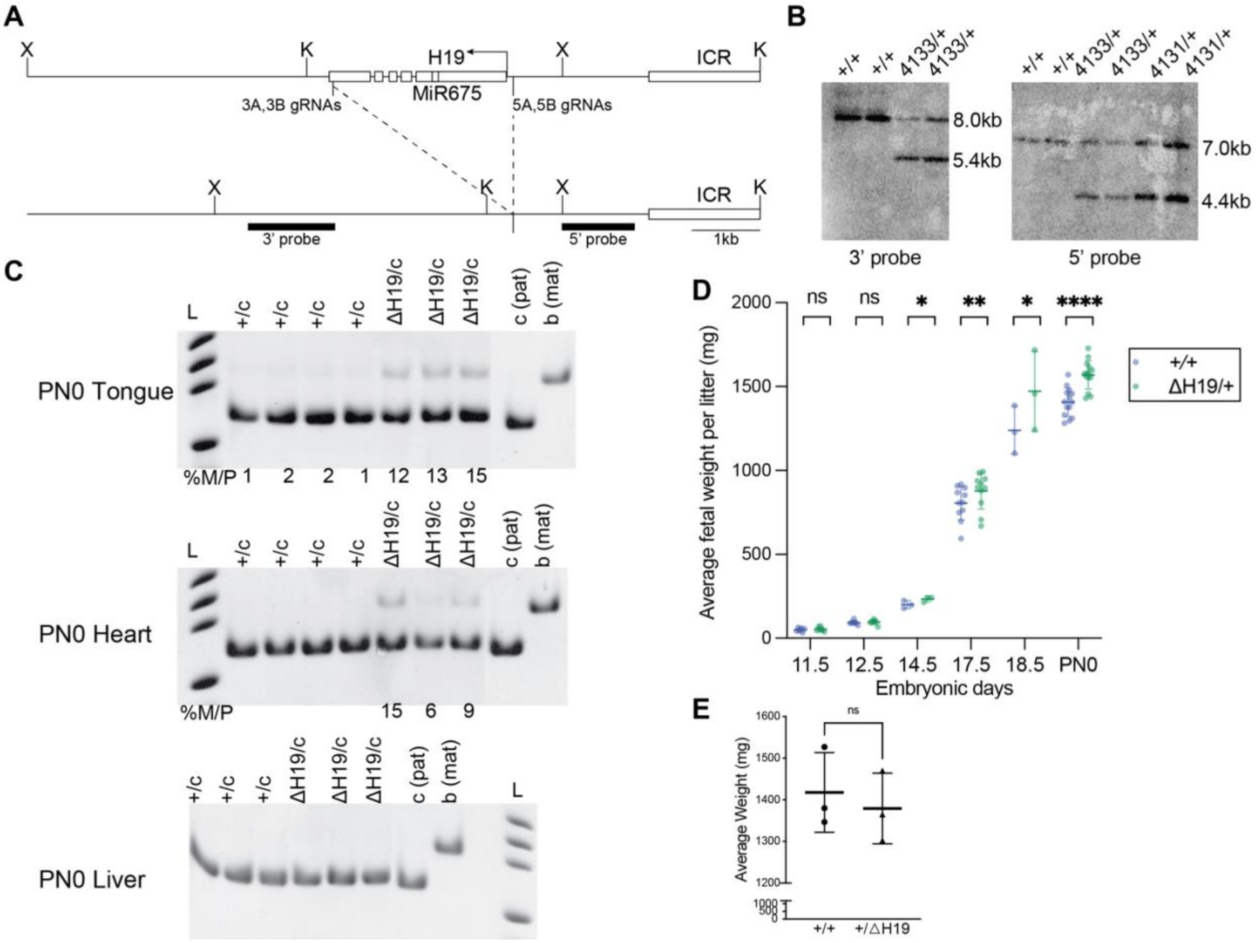


**Supplemental Figure 3. Characterization of *△H19* allele.**

(A) Targeting strategy to generate the *△H19* allele. The endogenous *H19* locus is shown with restriction sites, gRNA locations used to generate the deletion and binding sites for probes used in Southern blot analysis (thick lines). (B) Southern blot analysis of *△H19* allele. Founder 4131 line was generated using gRNA pair A, and founder 4133 line was generated using gRNA pair B (Supplemental Table 1). 3' probe and 5' probe were hybridized to XbaI- and KpnI-digested DNA, respectively. The sizes of the DNA fragments are shown on the right. (C) Allele-specific *Igf2* expression in wild-type and *△H19/+* neonatal tongue, heart and liver analyzed by restriction fragment length polymorphism (RFLP). Ladder, genotypes and c (*Mus castaneus*, paternal) and b (C57BL/6, maternal) allele controls are indicated above each gel. Quantification of band densitometry is shown below each gel, with percent of maternal allele expression relative to paternal allele indicated. No expression from the maternal *Igf2* allele was observed in liver. (D) Embryonic and neonatal body weight of the wild-type (blue) and *△H19/+* (green) samples at E11.5,

E12.5, E14.5, E17.5, E18.5 and PN0 (mean ± SD). 6 litters for E11.5, 6 litters for E12.5, 3 litters for E14.5, 11 litters for E17.5, 3 litters for E18.5,14 litters for PN0 are presented. (E) Body weight of +/*△H19* neonates (mean ± SD). 3 litters are presented. (D, E) Each data point represents an average weight of each genotype from one litter. Paired Student's t-test; **P* < 0.05, ***P* < 0.01, ****P* < 0.001, *****P* < 0.0001, ns = not significant.

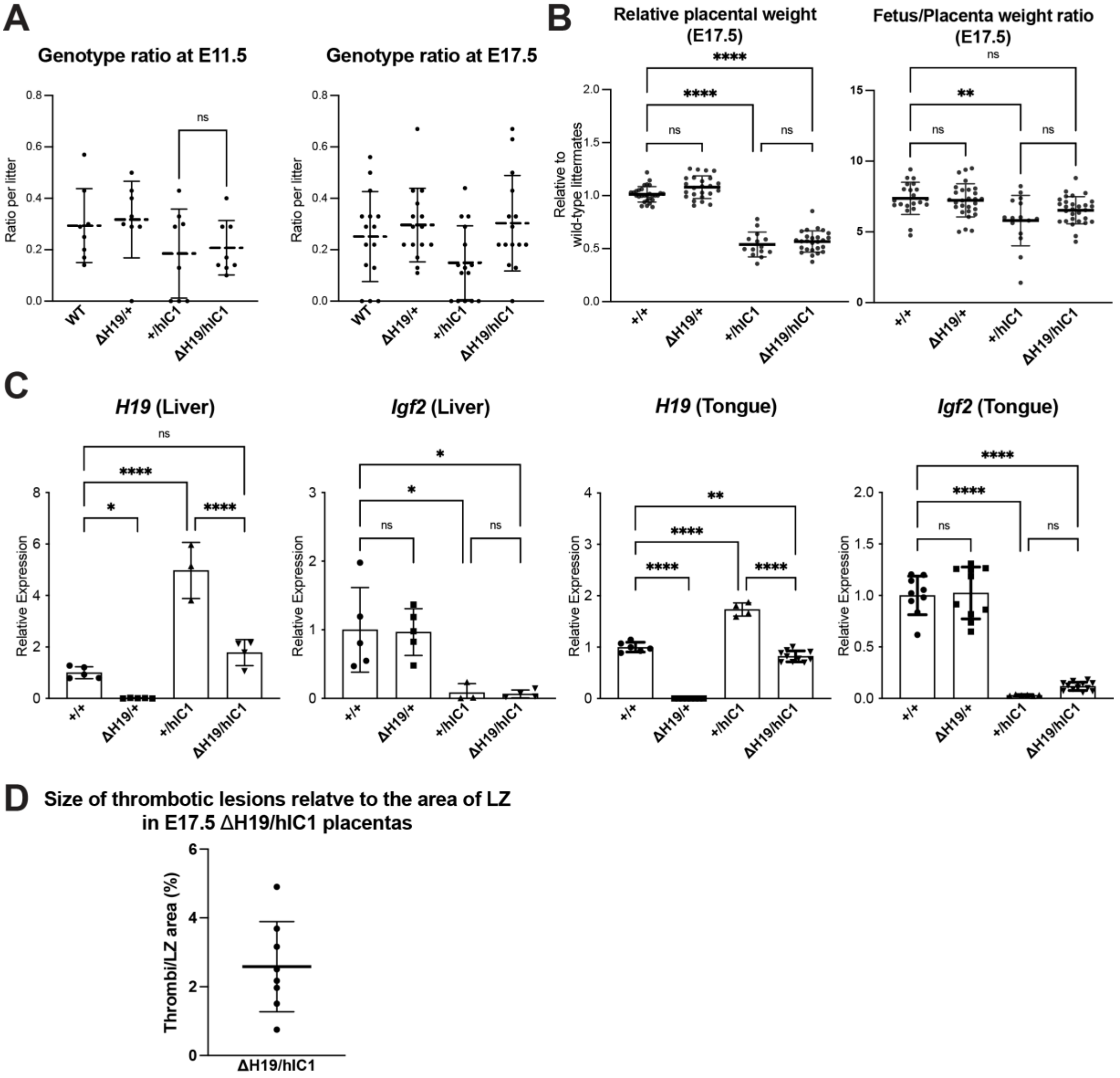


**Supplemental Figure 4. Supplementary data for rescue upon maternal *△H19* transmission.**

(A) Ratio of wild-type, *△H19/+, +/hIC1* and *△H19/hIC1* embryos observed in E11.5 and E17.5 litters (>5 pups) (mean ± SD). Each data point represents one litter. (B) (Left) Relative placental weights of E17.5 wild-type, *△H19/+, +/hIC1* and *△H19/hIC1* samples, normalized to the average placental weight of the wild-type littermates (mean ± SD). (Right) Fetal to placental weight ratio in E17.5 wild-type, *△H19/+, +/hIC1* and *△H19/hIC1* samples (mean ± SD). (C) Relative total expression of *H19* and *Igf2* in E17.5 wild-type, *△H19/+, +/hIC1* and *△H19/hIC1* liver and tongue samples (mean ± SD). (D) %Area of thrombotic clots in *△H19/hIC1* placentas, relative to labyrinth zone. (B, C, D) Each data point represents an individual conceptus from different litters. Statistics used are (A) Paired Student's t-test, (B, C) One-way ANOVA with Tukey's multiple comparisons test; **P* < 0.05, ***P* < 0.01, ****P* < 0.001, *****P* < 0.0001, ns = not significant.

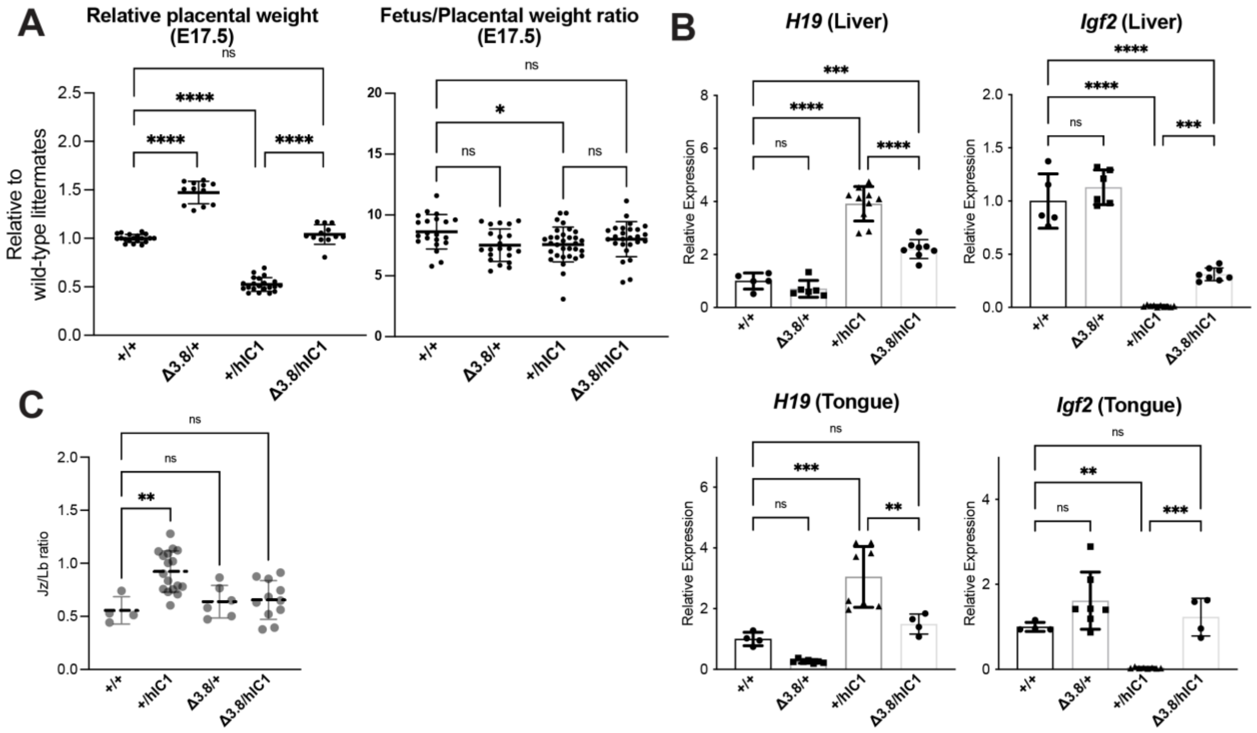


**Supplemental Figure 5. Supplementary data for rescue upon maternal *△3.8* transmission.**
(A) (Left) Relative placental weights of E17.5 wild-type, *△3.8/+, +/hIC1* and *△3.8/hIC1* samples, normalized to the average placental weight of the wild-type littermates (mean ± SD). (Right) Fetal to placental weight ratio in E17.5 wild-type, *△3.8/+, +/hIC1* and *△3.8/hIC1* samples (mean ± SD). (B) Relative total expression of *H19* and *Igf2* in E17.5 wild-type, *△3.8/+, +/hIC1* and *△3.8/hIC1* liver and tongue samples (mean ± SD). (C) Junctional zone (Jz) to labyrinth (Lb) ratio in E17.5 wild-type, *△3.8/+, +/hIC1* and *△3.8/hIC1* placentas. (A, B, C) Each data point represents an individual conceptus from different litters. One-way ANOVA with Tukey's multiple comparisons test; **P* < 0.05, ***P* < 0.01, ****P* < 0.001, *****P* < 0.0001, ns = not significant.

Chang_Supplemental Figure 6.

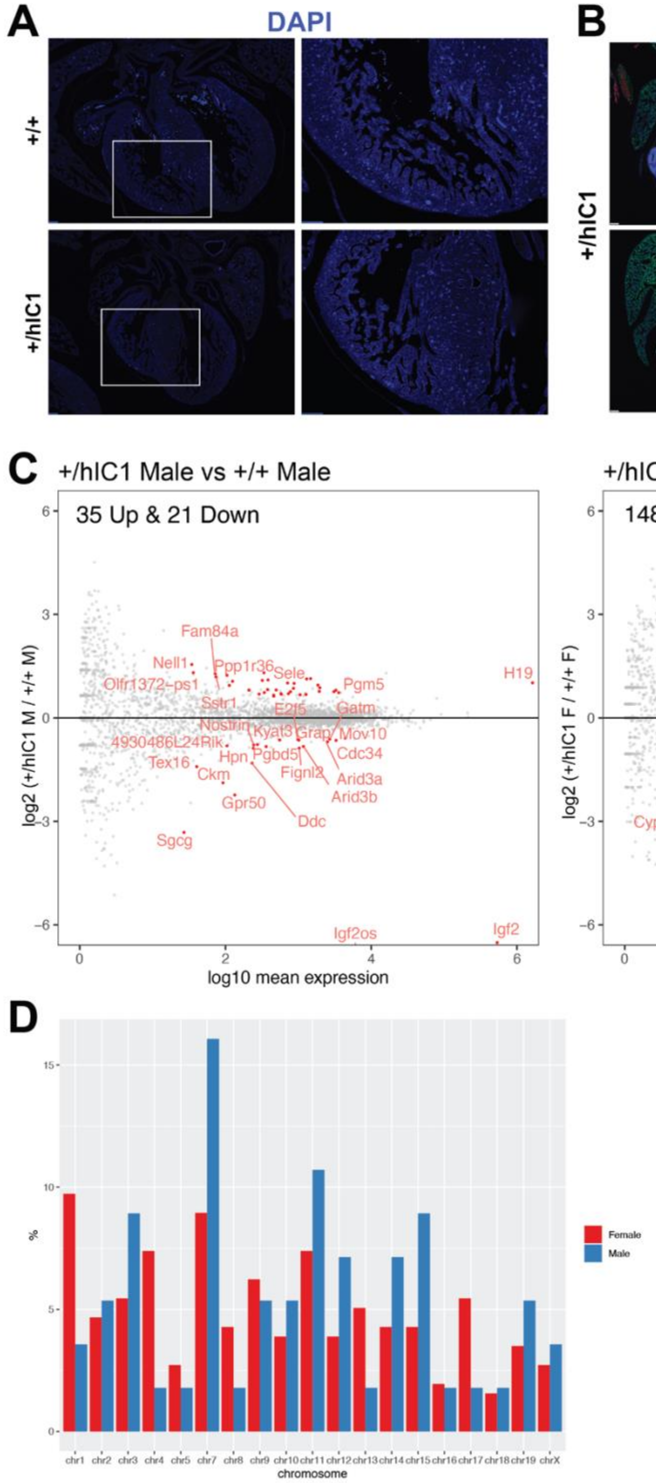


**Supplemental Figure 6. Supplementary data for differential gene expression of *+/hIC1* hearts.**

(A) DAPI (blue) on E17.5 wild-type and *+/hIC1* hearts from Figure 6C. Images on the right are enlarged from the boxed area of images on the left. Scale bars = 100 µm. (B) Immunofluorescence staining for MF20 (red), collagen I (green), and DAPI (blue) on E17.5 *+/hIC1* hearts. Images on the right are enlarged from the boxed area of images on the left. Scale Bars = 100 µm. (C) Same-sex comparisons between *+/hIC1* and the wild-type samples analyzed by RNA sequencing. (D) Chromosomal distribution of DEGs from comparisons between *+/hIC1* and wild-type males and females.

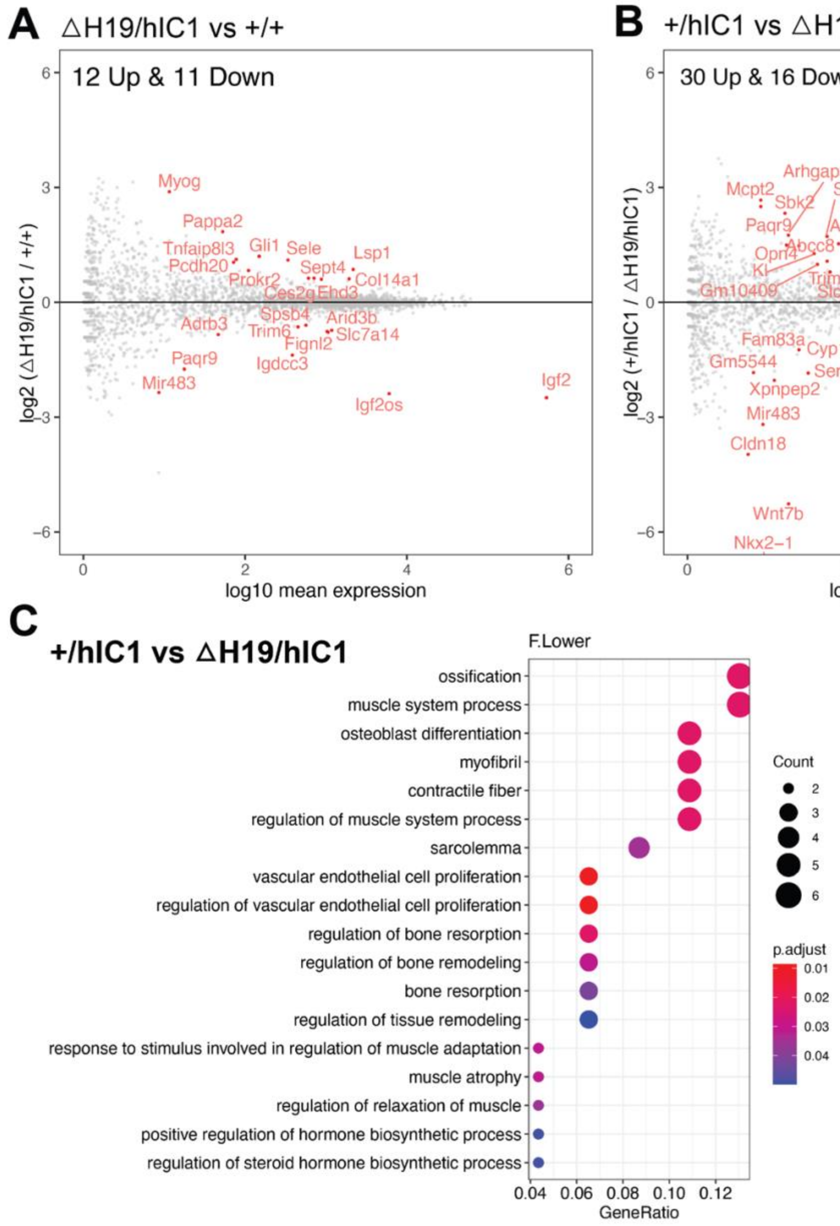


**Supplemental Figure 7.** (A) Volcano plot depicting comparison of *△H19/hIC1* and wild-type cardiac endothelial cells. (B) Volcano plot depicting comparison of *+/hIC1* and *△H19/hIC1* cardiac endothelial cells. (C) GO pathways that are enriched for 46 DEGs between *+/hIC1* and *△H19/hIC1* samples.

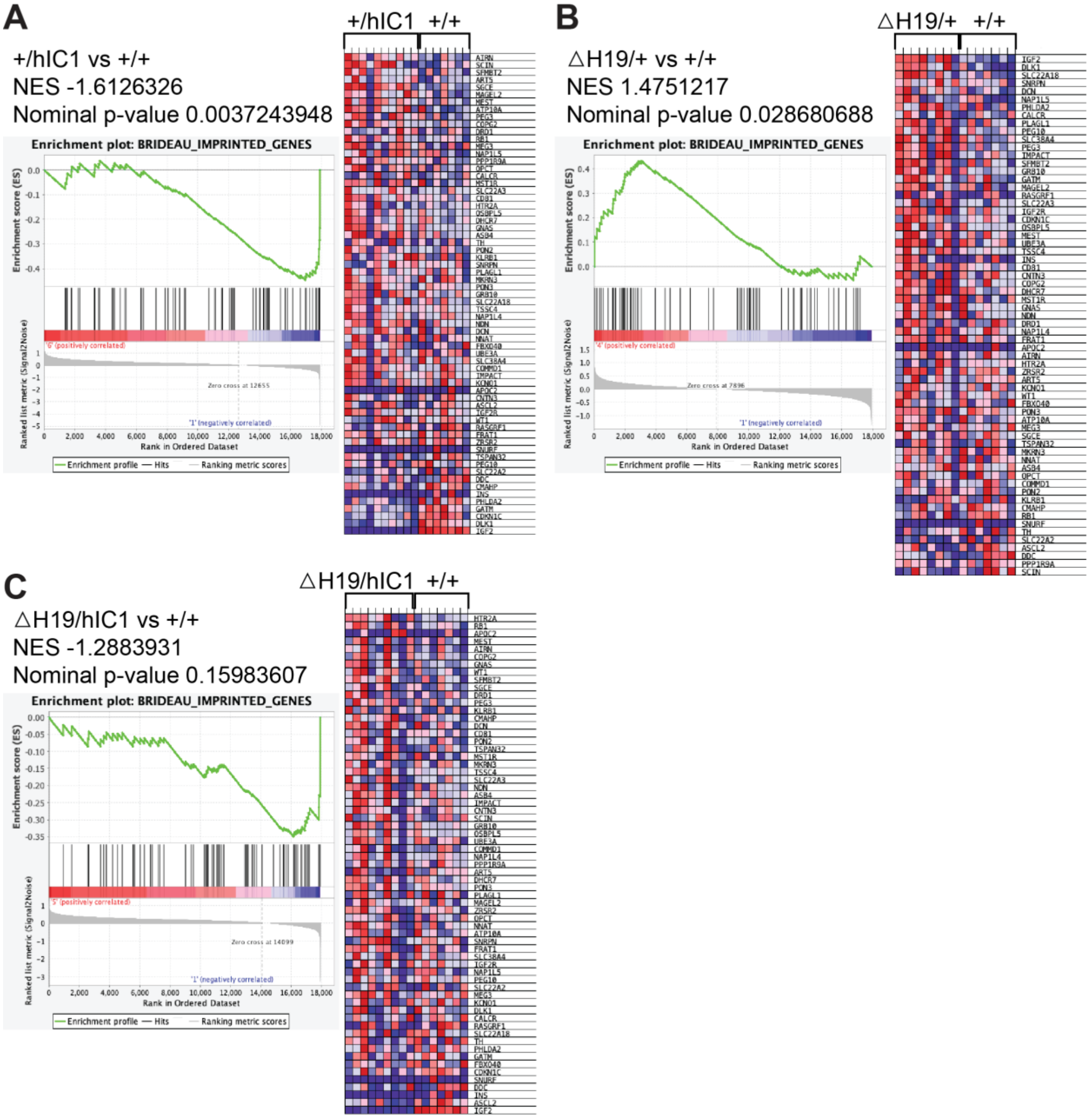


**Supplemental Figure 8. Enrichment tests for BRIDEAU_IMPRINTED_GENES using GSEA.** Normalized enrichment score (NES), nominal p-value, enrichment plot and expression heatmap of the analyzed gene set comparing (A) *+/hIC1* and wild-type, (B) *⊿H19/+* and wild-type, (C) *⊿H19/hIC1* and wild-type cardiac endothelial cells.

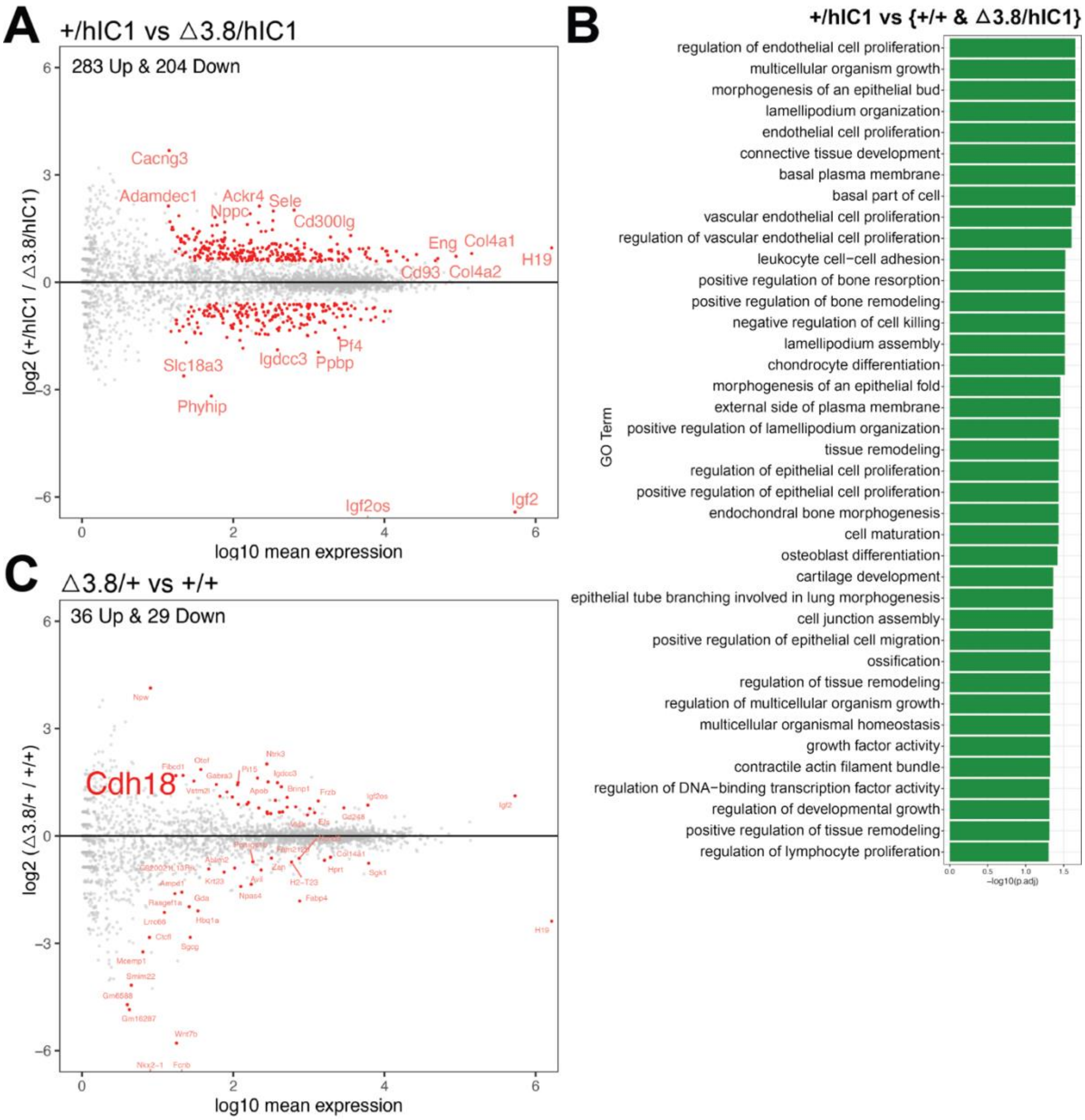


**Supplemental Figure 9.** (A) Volcano plot depicting comparison of *+/hIC1* and *△3.8/hIC1* cardiac endothelial cells. (B) GO pathways that are enriched for 116 DEGs that are commonly differentially expressed in *+/hIC1* compared to wild-type and *△3.8/hIC1* samples. (C) Volcano plot depicting comparison of *△3.8/+* and wild-type cardiac endothelial cells.

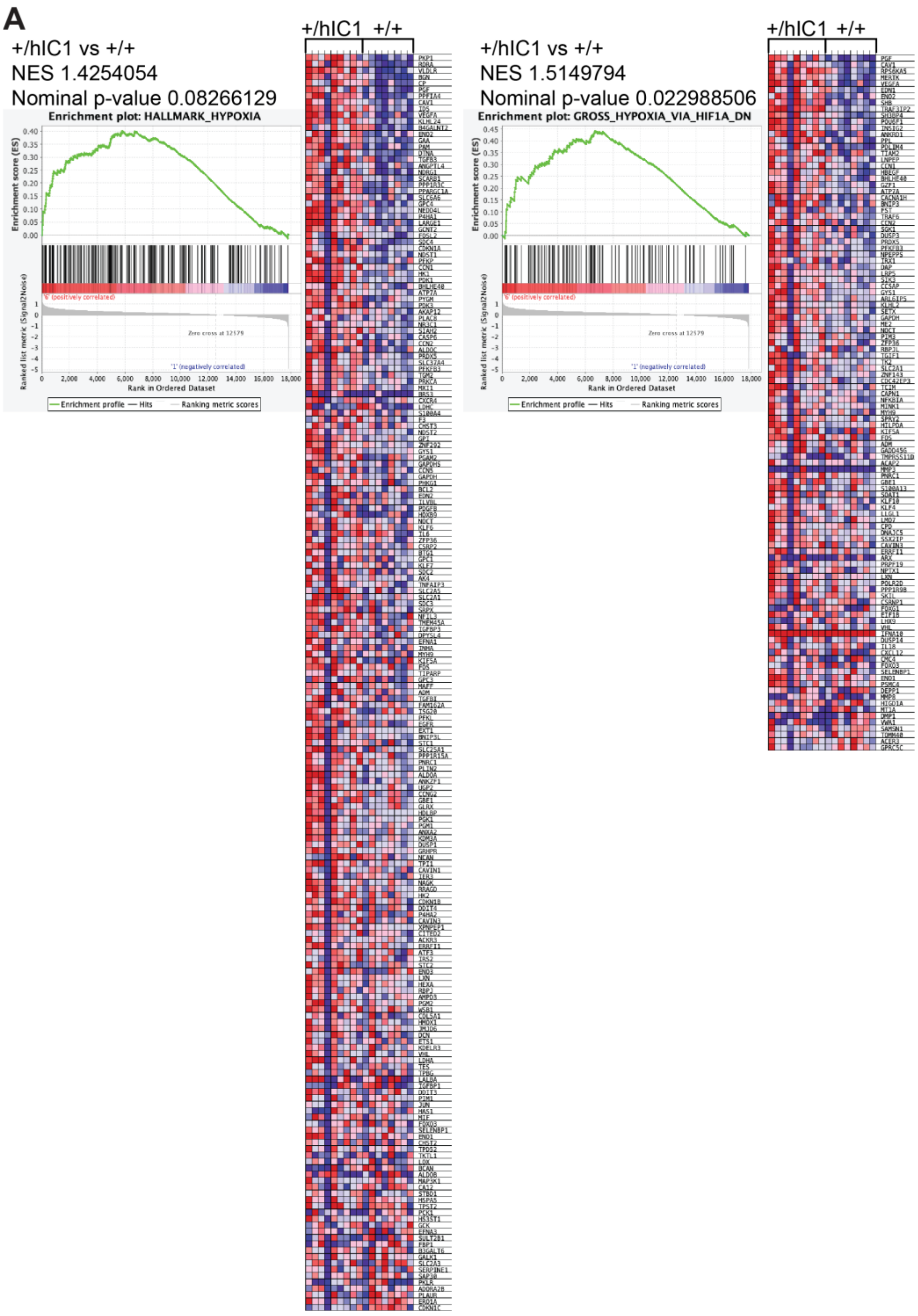


**Supplemental Figure 10. Enrichment tests for HALLMARK_HYPOXIA, GROSS_HYPOXIA_VIA_HIF1A_DN using GSEA.**

Normalized enrichment score (NES), nominal p-value, enrichment plot and expression heatmap of the (A) HALLMARK_HYPOXIA, (B) GROSS_HYPOXIA_VIA_HIF1A_DN gene set comparing *+/hIC1* and wild-type cardiac endothelial cells.
